# Peptide derived nanobody inhibits entry of SARS-CoV-2 variants

**DOI:** 10.1101/2022.04.21.489021

**Authors:** Nivya Mendon, Rayees Ganie, Shubham Kesarwani, Drisya Dileep, Sarika Sasi, Prakash Lama, Anchal Chandra, Minhajuddin Sirajuddin

## Abstract

Emergence of the new escape mutants of the SARS-CoV-2 virus has escalated its penetration among the human population and has reinstated its status as a global pandemic. Therefore, developing effective antiviral therapy against emerging SARS variants and other viruses in a short period of time becomes essential. Blocking the SARS-CoV-2 entry into human host cells by disrupting the spike glycoprotein-ACE2 interaction has been already exploited for vaccine development and monoclonal antibody therapy. Unlike the previous reports, our study used a 9 amino acid peptide from the receptor-binding motif (RBM) of Spike (S) protein as an epitope. We report the identification of an efficacious nanobody N1.2 that blocks the entry of pseudovirus containing SARS-CoV-2 spike as the surface glycoprotein. Moreover, we observe a more potent neutralizing effect against both the hCoV19 (Wuhan/WIV04/2019) and the Omicron (BA.1) pseudotyped spike virus with a bivalent version of the nanobody. In summary, our study presents a faster and efficient methodology to use peptide sequences from a protein-receptor interaction interface as epitopes for screening nanobodies against potential pathogenic targets. This approach can also be widely extended to target other viruses and pathogens in the future.

## Introduction

The severe acute respiratory syndrome coronavirus 2 (SARS-CoV-2) belongs to the betacoronavirus family and is the third member of the family after MERS-CoV and SARS-CoV that has infected the human population, resulting in moderate to severe pathogenic symptoms (1). With the frequent outbreaks of new variants, these numbers are increasing every day (2). Although several safe and effective anti-SARS-CoV-2 therapies have been recently developed which have been proven to prevent severe symptoms upon infection, none of them has been proven to be 100% effective. Partial vaccination, in turn, has put the SARS-CoV-2 virus under increased selection pressure resulting in a high mutation rate in the viral spike protein (3). With the emergence of novel variants and escape mutants of SARS-CoV-2, there is an urgent need to develop effective antiviral therapy that can single-handedly target both the current and emerging variants.

The surface of the SARS-CoV-2 virus is decorated with homotrimer Spike (S) envelope glycoprotein, the major antigenic determinant of the host immune response and one of the main coronavirus drug targets (4). The SARS-CoV-2 entry is orchestrated by the interaction of the receptor-binding domain (RBD) of its spike with the angiotensin-converting enzyme 2 (ACE2) receptor on the host cell membrane (5, 6). The precise contact is established by a set of conserved residues present in the receptor-binding motif (RBM) of the spike (7). Efforts have been made to prevent this first step of infection using antibodies or nanobodies that target the spike-ACE2 interaction (8–16). Several groups have also isolated antibodies directly from convalescent SARS-CoV-2 patients that showed promising neutralization against SARS-CoV-2 in-vitro and improved clinical outcomes in tested animals in vivo (17–22). Nanobodies, on the other hand, offer many advantages over conventional antibodies in terms of their size, high thermal stability, solubility, and ease of expression in bacteria hence making them easily scalable and cost-effective (23). Nanobodies, which are derived from the variable domain of the camelid heavy chain, retain specificity and affinity similar to conventional antibodies. Modularity in nanobodies allows oligomerization hence increasing their avidity and serum shelf life. (24). Nanobodies can be easily humanized which is critical for developing antiviral therapies for humans (25). Treatment with combinatorial nanobodies has been effective in neutralizing SARS-CoV-2 and preventing mutational escape (11). Many of such anti-SARS-CoV-2 therapies report the use of the entire spike protein or its RBD domain as the epitope to screen active antiviral antibodies. A limitation of using the entire spike protein or RBD as an epitope is that the isolated antibodies may bind to different regions of spike protein other than the ACE2 interacting domain, thus rendering them unspecific and inefficient in fully neutralizing the viral infection. Here, we report the isolation of a nanobody (N1.2) from a yeast nanobody display library using a small 9 amino acid peptide sequence as an epitope. The peptide was designed from the specific residues within the RBM showing high-affinity interaction with the spike protein with the ACE2 receptor. This nanobody, both monomer as well as tandem dimer interferes with ACE2 binding thus showing a potent virus neutralization activity in cellular models.

## Results

### Screening and identification of anti-spike nanobodies

The trimeric spike envelope protein of SARS-CoV-2 interacts with the cellular ACE2 (angiotensin-converting enzyme 2) receptor in the host cell plasma membrane to gain access to the cytoplasm (Figure 1A). We designed two peptide sequences from the spike envelope protein; peptide-1, residue 482-494, and peptide-2, residues 496 – 506 from the RBM of hCoV19 (Wuhan/WIV04/2019) (5). These peptides were used as bait (epitope) against the combinatorial yeast display library of nanobodies (26) and enrich potential nanobody sequences. The screening methodology involves two rounds of magnetic screening followed by fluorescent activated cell sorting (FACS) based enrichement of library (Figure 1B). From the FACS analysis, only peptide-1 showed positive enrichment of the nanobody clones (Figure 1B). Therefore, from here onwards we focused only on the nanobody population obtained from the peptide-1 screen. After FACS, the enriched yeast clones were individually analyzed using clonal selection and sequencing. Ten unique nanobody sequences were identified and purified, named N1.1 to N1.10 (Supplementary Figure 1 and Supplementary Table 1) (Methods).

**Figure 1:**
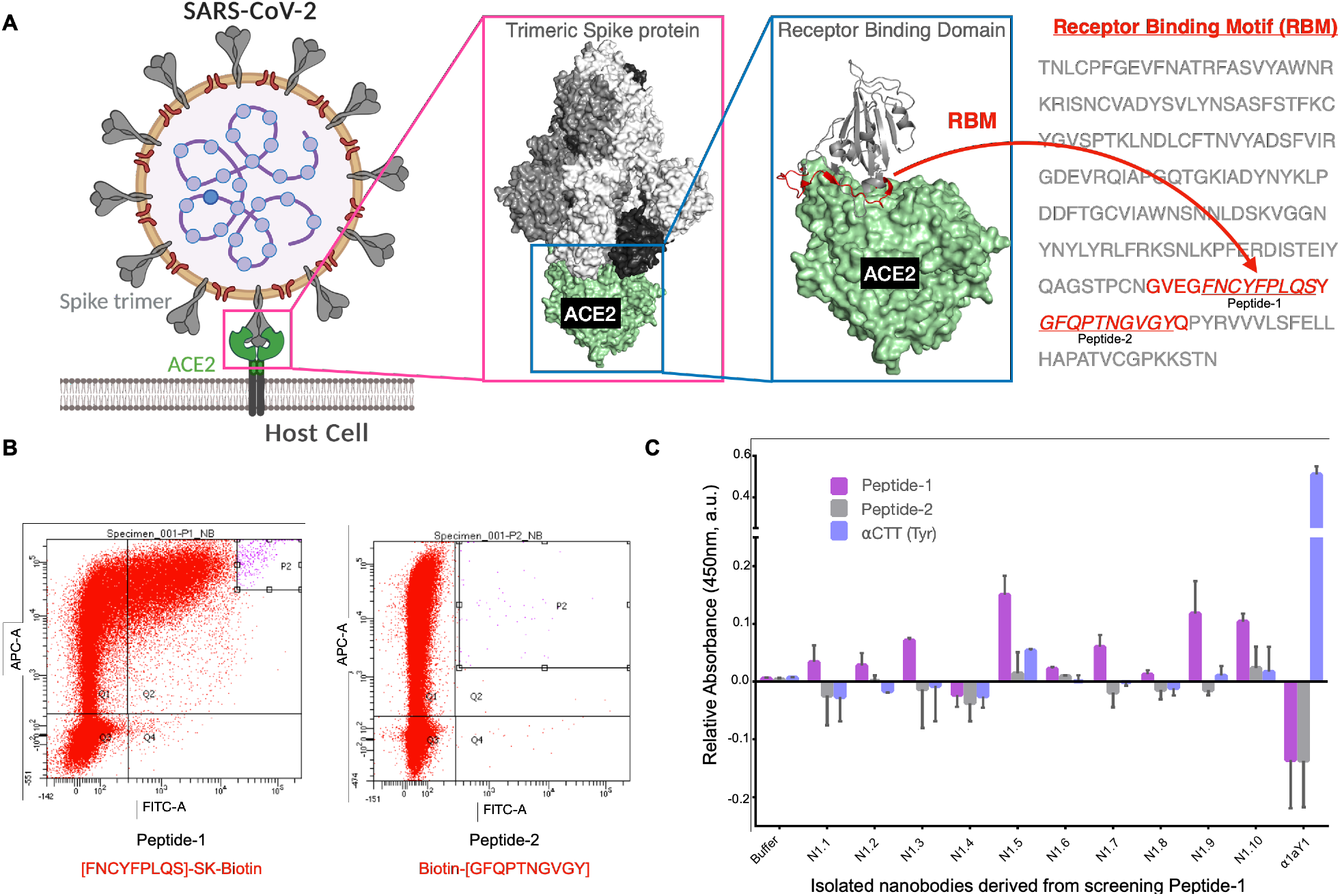
Identification of nanobodies against spike protein using peptides. **A**. Schematic illustration of SARS-CoV-2 virus entry into the host cell through ACE2 receptor. Cartoon model of molecular interaction between spike- and RBD-ACE2 derived from PDBID: 6VXX and 6LZG respectively. The Receptor Binding Motif (RBM) is highlighted in red in the ribbon model and sequence, the peptide-1 and -2 sequences are underscored and indicated. **B**. Representative images for peptide-1 and -2 sorting data on BD FACS Aria fusion cell sorter, the nanobody library labelled using c-myc marker in Alexa fluor-633 channel, and the peptides with streptavidin FITC. The P2 quadrant represents the double labelled yeast population indicating the extent of enrichment of yeast clones that has an affinity towards the respective peptide. **C**. ELISA assay of purified nanobodies against the peptide-1, -2, and tyrosinated alpha-tubulin CTT peptide. The A1aY1 protein against tyrosinated alpha-tubulin CTT peptide was included as an assay control.

The purified nanobodies were subjected to ELISA to characterise the interaction with peptide-1 and -2 (Methods). All the nanobodies showed good binding with the peptide-1 compared to the peptide-2, except N1.4 (Figure 1C). Suggesting that the enriched nanobody clones N1.1 to N1.10 are specific towards the peptide-1 epitope.

### Characterisation of nanobodies using flow cytometry based cellular assay

We then tested each of the purified nanobodies for their ability to block Spike: ACE2 interaction using a pseudovirus based neutralizing assay (Figure 2A). Pseudoviruses displaying spike envelope glycoprotein with a mCherry reporter were generated (27) (Methods). The viral infectivity was assessed using the mCherry expression, which was further quantified using fluorescence measurement (Methods).

**Figure 2:**
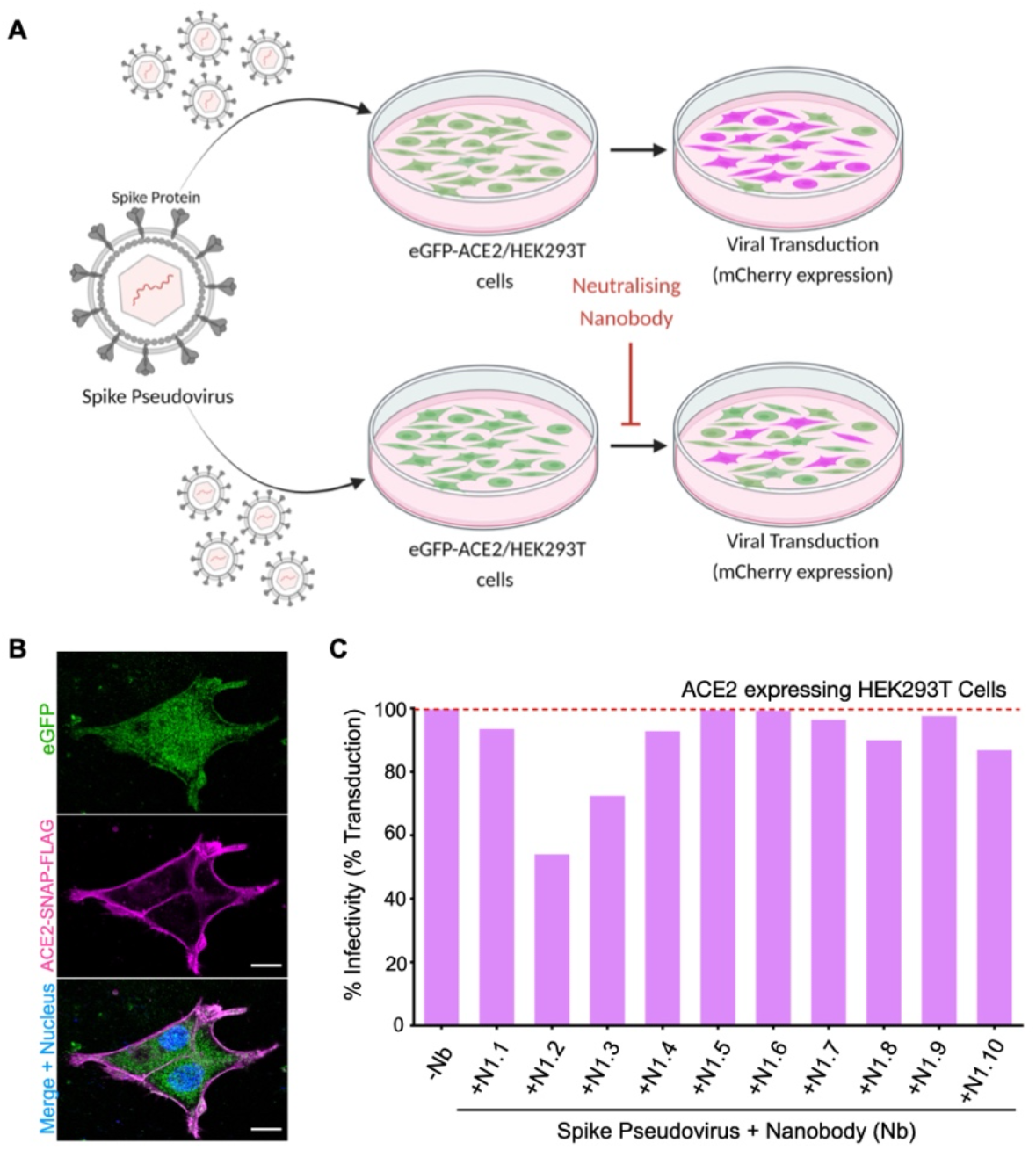
Characterization of nanobodies using virus entry assay. **A**. Cartoon representation of the virus entry assay using Spike pseudovirus. The pseudoviruses contain mCherry gene in their genome and upon entering the eGFP-ACE2/HEK293T cells the mCherry will be expressed and the fluorescence can be used for quantification of the virus infection. **B**. Maximum intensity Z-projection of confocal images obtained from eGFP-ACE2/HEK293T (magenta), eGFP (green), and nucleus (blue). **C** FACS based quantification of spike pseudovirus infection into eGFP-ACE2/HEK293T (magenta) in the presence of various nanobodies as indicated (Methods). The percentage of infectivity is normalized to the percentage of mCherry positive cells in the absence of nanobody in the assay. For raw FACS data and details regarding the assay refer to Supplementary Figure 4 and Methods.

Since the ACE2 expression in HEK293T cells is lower compared to other cell lines (28, 29), we generated an ACE2 stable expressing HEK293T cells, herein referred to as eGFP-ACE2/HEK293T cells (Methods). The eGFP-ACE2/HEK293T stable cell lines also co-expressed eGFP as a fluorescent marker to facilitate identifying ACE2 expressing cells (Figure 2B and Supplementary Figure 2) (Methods). The eGFP-ACE2/HEK293T cells were transduced with the varying dilutions of spike pseudoviruses (Methods). Quantification using FACS analysis showed ∼75%, 55%, 16%, and 4% of cells transduced with mCherry fluorescence correlating with the varying dilution of pseudoviruses respectively (Supplementary Figure 3A and 3B). We characterized the transduction efficiency of the pseudovirus as a function of its multiplicity of infection (MOI). In order to test the efficacy of nanobodies towards blocking the viral entry, we choose to perform experiments with pseudovirus MOI that yield >70% transduction efficiency and low cellular cytotoxicity.

25µM of each nanobody, N1.1 to N1.10 was preincubated with the Spike pseudovirus before infecting eGFP-ACE2/HEK293T cells (Methods). Except for N1.2, and N1.3, all the nanobodies showed mCherry fluorescence (i.e., no effect on pseudovirus entry) upon transduction in the eGFP-ACE2/HEK293T cells (Figure 2C and Supplementary Figure 4). Interestingly, the N1.2 nanobody showed maximum efficiency in neutralizing the Spike pseudoviruses compared to the remainder of the nanobodies (Figure 2C). Therefore, we decided to further characterize the N1.2 nanobody as a potential nanobody against Sars-CoV2 virus entry.

### Microscopy based cellular assay to assess N1.2

In our virus titration assays, we noticed that the FACS-based quantification does not scale according to the mCherry expression (Supplementary Figure 3A and 3B). Therefore, we simultaneously correlated the mCherry fluorescence versus virus infection using confocal microscopy-based quantification (Methods). In comparison to FACS data, the confocal microscopy based measurement, showed a sharp decline in mCherry fluorescence, i.e., >80% decrease in mCherry fluorescence when 1/10^th^ of the virus dilution was used for infection (Supplementary Figure 3C-E). Therefore, hereafter, we used confocal microscopy based measurement to assess the neutralizing potential of N1.2 nanobody against spike pseudovirus. In the pseudoviral assay, we used different concentrations of N1.2 (5µM, 10µM, and 25µM) and could observe a linear reduction in the infectivity of the spike pseudoviruses towards the eGFP-ACE2/HEK293T cells (Figure 3B, C and Supplementary Figure 5). To achieve a more potent inhibitory effect with N1.2, we engineered and purified a bivalent nanobody, in which the N1.2 sequence was placed in tandem separated by a glycine-serine linker (Methods) (Supplementary Figure 1 and Supplementary Table 1). The tandem N1.2, termed (N1.2)_2_ showed a significant reduction in pseudoviral transduction at 5µM and 10µM concentrations (Figure 3B and C), suggesting a cooperative effect in neutralizing the spike pseudovirus.

**Figure 3:**
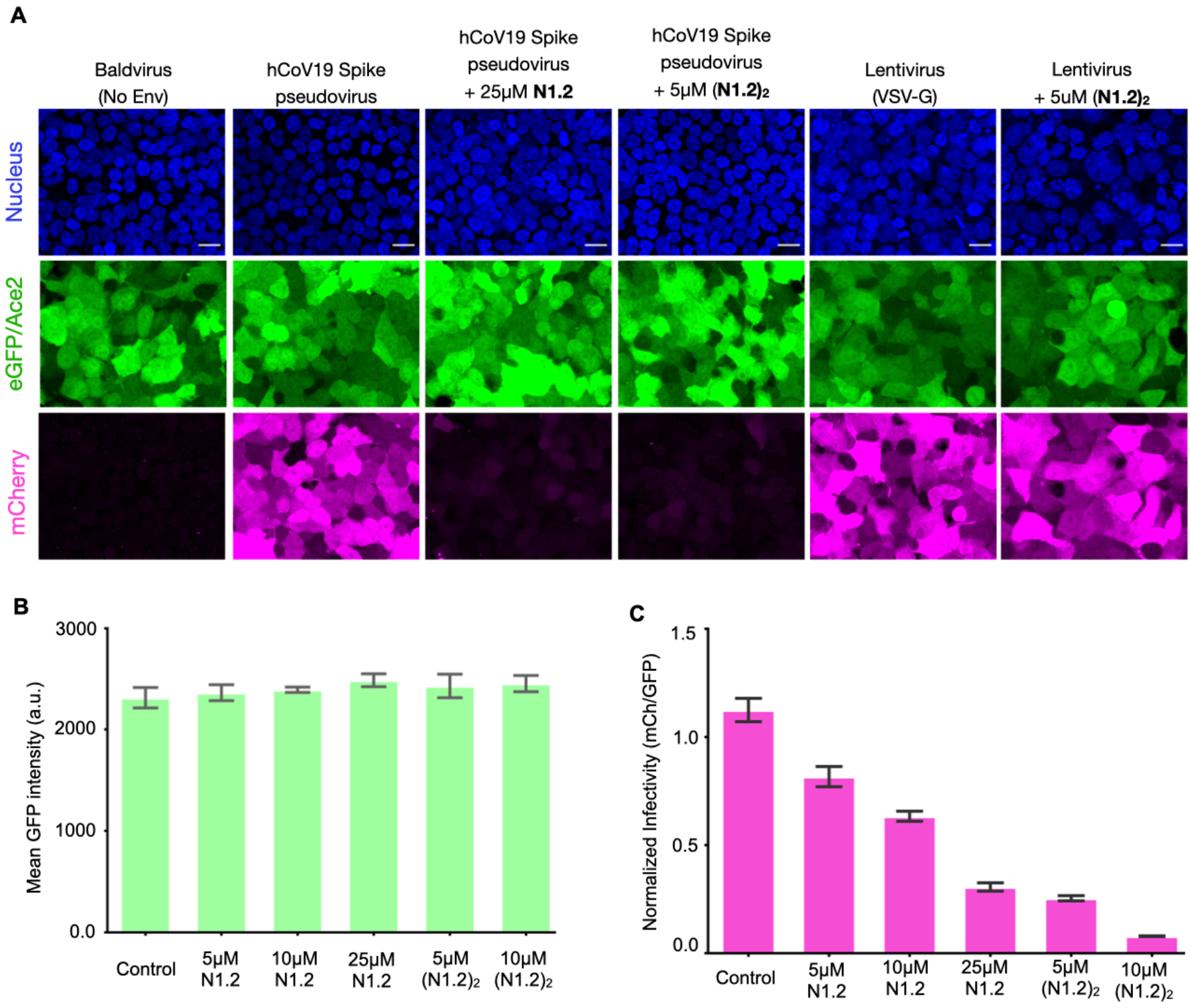
Assessing the efficacy of nanobody N1.2 using confocal microscopy. **A**. Maximum intensity Z-projection of confocal images of bald and spike pseudovirus in the absence and presence of various concentrations of monovalent and bivalent nanobody N1.2 as indicated. The nucleus is shown in blue, the eGFP fluorescence (in green) is a proxy for ACE2 expression in HEK293T cells, and mCherry (in magenta) indicates the extent of virus entry into the eGFP-ACE2/HEK293T cells. Scale bar = 20 microns. **B**. Mean eGFP fluorescence from individual experiments as indicated, control group represents spike pseudovirus assay in the absence of nanobody. **C**. Normalized infectivity quantified from mCherry over eGFP fluorescence for experiments with monovalent and bivalent nanobody N1.2 as indicated. For both B and C mean was calculated from three individual experiments and error bars are the standard error of the mean.

To confirm if this inhibitory effect of N1.2 and (N1.2)_2_ is achieved by its direct binding to spike envelope protein of SARS-CoV-2, we performed control experiments using VSV-G pseudoviruses and bald pseudoviruses (no glycoprotein envelope control) (Figure 3A and Supplementary Figure 5). These control VSV-G pseudoviruses transduce the stable cells similar to the spike pseudoviruses in the presence and absence of nanobody N1.2 or (N1.2)_2_ (Figure 3A. This indicates that the neutralizing effect of N1.2 and (N1.2)_2_ is obtained by direct binding of the nanobody to the spike protein. Therefore, the engineered nanobody N1.2 can be effectively applied for antiviral therapy against Covid-19.

### Efficacy of N1.2 against Omicron spike variant

We also tested the efficacy of the N1.2 against the newly emergent Omicron variant, using the Omicron spike containing pseudoviruses expressing mCherry (Methods). First, we confirmed the infectivity of varying concentrations of Omicron spike pseudovirus against eGFP-ACE2/HEK293T cells and determined an effective MOI (Supplementary Figure 3A and 3B). Similar to the hCoV19 (Wuhan/WIV04/2019), the mCherry fluorescence was also scaled with the virus MOI for the Omicron variant spike pseudovirus in the confocal microscopy-based mCherry quantification (Supplementary Figure 3C, D and E). We further tested the tandem nanobody (N1.2)_2_ neutralizing effect against the Omicron spike pseudovirus in GFP-HEK293T and eGFP-ACE2/HEK293T (Methods). The hCoV19 (Wuhan/WIV04/2019) and Omicron spike pseudovirus were efficiently blocked by 5uM of (N1.2)_2_ as observed from the confocal images in both cell lines (Figure 4A and Supplementary Figure 6). Quantification of infectivity showed that in the presence of 5µM of (N1.2)_2_ the infectivity drops to <10% (Figures 4B and 4C). Further, indicating the broad neutralizing ability of the N1.2 nanobody reported here against the current SARS-CoV2 Omicron variant.

**Figure 4:**
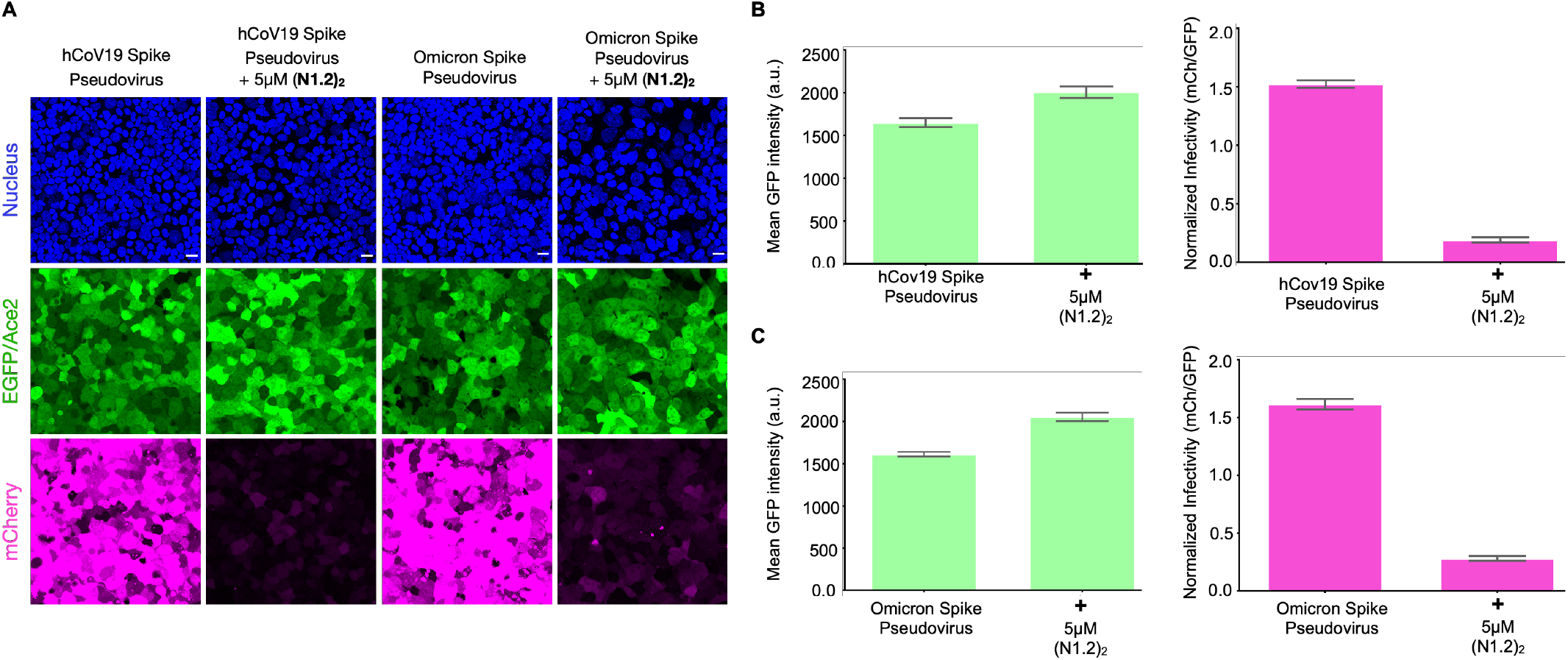
Comparison of nanobody N1.2 against hCoV19 and Omicron spike variant. **A**. Maximum intensity Z-projection of confocal images of hCoV19 and Omicron spike pseudovirus in the presence of 5uM bivalent nanobody N1.2 as indicated. The nucleus is shown in blue, the eGFP in green, and mCherry in magenta. Scale bar = 20 microns. **B & C**. Mean eGFP (green column graphs) and normalized infectivity quantification of hCoV19 and Omicron spike (magenta column graphs) in the absence and presence of 5uM bivalent nanobody (N1.2)_2_ as indicated. N = 3 experiments and error bars = s.e.m.

## Discussion

Since 2020 the SARS-CoV2 virus has evolved into many subtypes and variants that are still prevalent across the globe (30). Therefore it is pivotal to identify broad-spectrum therapeutics that can effectively work against all SARS-CoV2 subtypes and variants. Equally important is to establish validated pipelines that can expedite identifying new therapeutics against emerging SARS-CoV2 variants or new viruses. In this study, we have addressed both the needs: first, we have identified a broad spectrum nanobody that neutralizes the original Wuhan (WIV04) and Omicron spike pseudovirus. Secondly, we have achieved the nanobody identification using a peptide as antigen, unlike the whole spike or RBD protein of SARS-CoV2 described earlier (13, 16, 31–33). The nanobodies or antibodies obtained by using a whole spike or RBD bind to different regions of the spike protein and only a fraction of them were effective in neutralising the virus (33). Largely because these antibodies target the regions of the spike protein other than the ones involved in spike-ACE2 interaction. Using peptides as antigens has been widely used for generating potent antibodies targeting the desired epitope, including the recent example for SARS-CoV-2 antibodies (34). However, peptides and flexible loops from a target protein have seldom been used as antigenic baits for screening display libraries. We previously reported a nanobody against tubulin post-translation modification using the flexible carboxy-terminal tail peptide (35). Applying a similar principle, here we show that a 9 amino acid peptide from the receptor-binding motif (RBM) of spike protein can also be employed as an epitope against a nanobody display library. This approach of using synthetic peptides as an epitope, instead of purified whole protein molecules will certainly accelerate and fine tune the entire screening process and yield nanobodies/antibodies that are efficacious in neutralizing the future potential targets /viruses.

The nanobody N1.2 described in this study is effective against both the spike proteins from the original hCoV19 (Wuhan/WIV04/2019) and the recent Omicron variant. Sequence alignment of the peptide-1 and its surrounding region that was used in screening the nanobody shows the high similarity between the variants (Supplementary Figure 1). Suggesting that epitopes from the conserved RBM region such as the peptide-1 can yield specific nanobodies or antibodies against the SARS-CoV-2 virus, yet possess broad neutralizing ability against the existing and emerging variants. Almost all the SARS-CoV2 variants reported to date utilize ACE2 receptor binding for the host cell entry and the RBM region of spike protein predominantly contributes to this interaction (36, 37). Therefore, we anticipate that the nanobody N1.2 will remain effective in neutralizing the emerging variants as long as the SARS-CoV2 virus uses ACE2 for gaining entry into the host cells. Additionally, the nanobodies offer advantages in engineering the valency and thereby increasing the efficacy in neutralization against the target molecules. Indeed when we engineered the nanobody N1.2 into a bivalent version, (N1.2)_2_ we observed a cooperative effect in neutralizing the pseudovirus even at the highest concentrations. Together, our study offers a platform to identify broad spectrum nanobodies specific against SARS-Cov2 and its variants, which can be exploited for other therapeutically relevant targets.

## Materials and Methods

### Peptide epitopes for screening

The peptide sequences were derived from two different regions of the receptor-binding motif (RBM) of spike proteins that have been crucial for the interaction of viral spike protein with the cellular ACE2 receptor. The following peptide sequences from the receptor-binding domain (RBD) of the spike were synthesized from LifeTein with a biotin tag:

Peptide-1: [FNCYFPLQS]S-K-Biotin

Peptide-2: Biotin-[GFQPTNGVGY]

Sequence Alignment for Covid Variants

The hCoV19 spike (Wuhan/WIV04/2019), GISAID (EPI_ISL_402124) construct is a kind gift from Prof. Nevan Krogan, UCSF, USA (38). The sequence used for alignment was obtained from NCBI Genbank. The accession number for Alpha (B1.1.7): MW487270.1, Delta (B1.617.2): OK091006.1, and BA.2: OM296922.1. The Omicron MC_0101274 was purchased from Genscript.

### Screening yeast-display nanobody library of nanobodies

A combinatorial yeast-display library of nanobodies (NbLib) was obtained from Kerafast, Inc. The estimated diversity of the library is around 5*10^8 unique nanobody clones expressed onto the surface of the yeast cells. The detailed protocol for screening nanobodies against the peptides has been adopted from the method described earlier (26). The frozen vial of NbLib (5-fold excess of library diversity) was grown in 1L Yglc4.5-Trp media (3.8g/L of - Trp drop-out media supplement, 6.7g/L yeast nitrogen base, 10.4g/L sodium citrate, 7.4g/L citric acid monohydrate, 20g/L glucose and 10ml/L PenStrep, pH4.5) at 30°C for 24-48hr and passaged thrice before the screening. The freshly passaged NbLib (5*10^9 cells, 10-fold excess of library) was induced in galactose containing media (-Trp +galactose; 3.8g/L of -Trp drop-out media supplement, 6.7g/L yeast nitrogen base, 20g/L galactose and 10ml/L PenStrep, pH6.0) at 20°C for 72hr to achieve sufficient expression in the library.

#### Magnetic selection (MACS)

Around 5*10^9 nanobody expressing yeast cells were pelleted to remove media and washed with 10ml selection buffer (20mM HEPES pH 7.5, 150 mM sodium chloride, 0.1% (w/v) bovine serum albumin, and 5 mM maltose). The cells were resuspended in 4.5ml of selection buffer. First, we performed negative magnetic selection by incubating the above-resuspended cells with 200μl of anti-biotin microbeads (Miltenyi Biotec, catalogue no. 130-090-485) and 200μl of streptavidin microbeads (Miltenyi Biotec, catalogue no. 130-048-102)at 4°C for 1hr. Post-incubation the cells were pelleted, resuspended in 5ml selection buffer, and were allowed to pass through (via gravity flow) the LD column (Miltenyi Biotec, catalogue no. 130-042-901) placed on Miltenyi MACS magnet (using Midi MACS separator) pre-equilibrated with 5ml selection buffer. The unbound cells were collected and the column was washed with an additional 2ml of the selection buffer to flush out the remaining cells from the column. The cells were pelleted and resuspended in a 3ml selection buffer for positive selection with the peptides.

##### First MACS selection

The cells were incubated with a 10μM concentration of peptide-1/peptide-2 at 4°C for 1hr, following which 100μl of anti-biotin microbeads were added and incubated for an additional 20min at 4°C. The cells were pelleted and washed with a 3ml selection buffer. The cells were resuspended in a 3ml selection buffer and passed through the pre-equilibrated (with 5ml selection buffer) LS column (Miltenyi Biotec, catalogue no. 130-042-401) placed on the Miltenyi MACS magnet. The LS column was washed with an additional 8ml of the selection buffer to remove unbound cells from the column. The column was removed from the magnet and added 3ml of the selection buffer to the column. Using a plunger eluted all the bound yeast cells from the column in 50ml falcon. The cells were pelleted and resuspended in -Trp +glucose media (3.8g/L of -Trp drop-out media supplement, 6.7g/L yeast nitrogen base, 20g/L glucose, and 10ml/L PenStrep, pH6.0) for growth at 30°C for 48hr.

##### Second MACS selection

Took 10^9 first magnetic sorted cells and induced them in -Trp +galactose media for 72hr. Around 10^8 freshly induced yeast cells were taken and pelleted. The cells were washed with a 5ml selection buffer and resuspended in a 3ml selection buffer. Added 10μM of the respective peptide and incubated at 4°C for 1hr. To this added 100μl of streptavidin microbeads and incubated for an additional 20min at 4°C. The cells were pelleted, washed with 3ml selection buffer, and resuspended in 3ml of selection buffer to pass them through the pre-equilibrated LS column placed on the Miltenyi MACS magnet. Washed the column with 8ml additional selection buffer, followed which the column was removed from the magnet and plunged all the bead-bound cells from the column by 3ml selection buffer. The cells were pelleted and resuspended in 5ml -Trp +glucose media and kept for growth at 30°C for 48hr.

#### Fluorescent Activated Cell Sorting (FACS)

Finally, the second MACS sorted culture was induced (around 10^9 cells) in galactose media at 20°C for 72hr and performed cell sorting experiments to enrich the high-affinity nanobody against the target peptides. Here, we took 10^7 cells and pelleted them in a microcentrifuge vial. These cells were washed with 1ml selection buffer and resuspended the cells in 100μl selection buffer. The cells were incubated with 100μM of the respective peptide and 1:200 dilution of rabbit anti-HA tag antibody (Sigma; catalogue no. Cat # H6908) at 4°C for 1hr. Cells were washed with 1ml selection buffer and resuspended in 100ul of selection buffer for secondary antibody staining. Cells were incubated with 1:200 dilution of goat anti-rabbit Alexa fluor-647 antibody (Invitrogen) and 1:100 dilution of neutravidin fluorescein conjugate (Invitrogen; FITC, catalogue no. A2662, used for the first FACS) at 4°C for 30min. Post-incubation cells were washed twice with a 1ml selection buffer to remove unbound reagents. Cells were resuspended in 1ml selection buffer just before sorting them on BD FACS Aria Fusion for double-positive cells, keeping unstained and single stained controls (Central Imaging and Flow Cytometry Facility at the National Centre for Biological Sciences [NCBS]). 0.1–1% of the cells, positive for both the fluorophores (for Neutravidin-FITC and Alexa Fluor 647 fluorophores) which are represented as the P2 population in the Q2 quadrant of the sorting layout. A total of around 8000-10,000 cells were collected and grown in 5 ml fresh -Trp +glucose media at 30°C for 48hr. The freshly grown cells were propagated in larger volumes in the same glucose media (250–500 ml) to make stocks (-Trp +glucose media and 10% DMSO) and further characterising individual clones for their affinity with the respective peptide. After the sorting experiment, the cells were plated onto an agar plate (-Trp +glucose +15 g/liter agar), and 10 single yeast colonies of post-sorted cultures were analysed on a flow cytometer (Thermo, Attune) for their binding with the respective peptide. Only peptide-1 screening yielded nanobody clones from the library having sufficient binding affinity for peptide-1 therefore, we have only focused to characterise nanobodies obtained from peptide-1 screening. We have isolated plasmids from these 10 individual yeast clones from peptide-1 screening to identify the protein-coding sequences of these nanobodies for their biophysical and cellular validation.

### Cloning and protein purification

The nanobody gene was amplified from isolated yeast colonies and cloned between HindIII and XhoI sites in a pET-22b(+) plasmid containing a C-terminal 6x histidine tag. The N1.2 nanobody sequence was first cloned with Linker AS-(G_4_S)_3_-G in a CMV vector (backbone from Addgene # 12298) using 5’-tccggtggcggaggctccggtggcggaggttccggacaggtgcagctgcaggaaagcgg-3’ and 5’-cacactggatcagttatctatgcggccgctcagtggtggtggtggtggtgctcgaggc -3’ primers. The sequence of N1.2 along with the AS(G_4_S)_3_G linker was amplified using 5’-gccagggcacccaggtgaccgtgagcagcgctagcggtggtggaggctccggtggcggaggc-3’ and 5’-cagccggatctcagtggtggtggtggtggtgctcgaggctgctcacggtcacctgggtgccc-3’ primers. This amplified construct of linker-N1.2 was cloned in pet22B amplified vector (5’-cggagcctccaccaccgctagcgctgctcacggtcacctgggtgccc-3’ and 5’-tcgagcaccaccaccaccaccactgagatccggctgctaac-3’) already cloned with N1.2 sequence (between NcoI and XhoI) using Gibson assembly method. The resulting construct contains an amino-terminus pelB sequence, two tandem sequences of N1.2 spaced with AS(G_4_S)_3_G linker, and a C-terminus 6X-Histidine tag.

The nanobodies were purified from Rosetta (DE3) cells in an LB medium by inducing at 0.5 OD with 0.5mM IPTG at 20 □C. After overnight induction, the cells were pelleted down and the protein was extracted by osmotic shock by resuspending in 0.5□M sucrose, 0.2□M Tris, pH 8, 0.5□mM EDTA followed by water in a 1:3 ratio with 1hr of stirring at 4°C. The lysate was adjusted to contain 150□mM NaCl, 2□mM MgCl_2_, and 20□mM imidazole and was subjected to centrifugation at 18000 RPM at 4°C to remove cell debris. The supernatant was loaded on 5ml HisTrap HP (GE Healthcare). The column is subsequently washed by high salt buffer (20mM Tris pH8, 500mM NaCl) and low imidazole buffer (20mM Tris pH 8, 100mM NaCl, 100mM imidazole pH 8). The protein is eluted with 20mM Tris,100mM NaCl, 400mM imidazole pH 8. The eluted protein is concentrated in 3KDa cut-off centricon (UFC9003) and buffer exchanged (20mM Tris pH 8, 100mM NaCl). The protein was aliquoted, flash-frozen, and stored at -80 □C. (26).

### Cell culture experiments

Wild type mammalian HEK293T cells and LentiX-293T cells (Takara Bio, catalogue no. 632180) were used in this study for pseudotyped spike virus production and viral transduction assay. Both these cell lines were grown in DMEM (ThermoFisher Scientific; catalogue no. 11995065) media supplemented with 10% FBS, 1× PenStrep (Gibco, Thermo Fisher Scientific; catalogue no. 15–140-122), and 1× GlutaMAX (Gibco, Catalog number: 35050061) in a humidified 37°C incubator with 5% CO_2_ supply.

### Spike pseudotyped virus production

Freshly passaged HEK293T cells were seeded in a 100mm cell culture plate and grown-up to 70-80% confluency. The media of the cells were changed to 10 ml complete DMEM without PenStrep before transfection. For viral particle production, 5 µg pHR lentiviral vector cloned with mCherry fluorescent protein, 3.75 µg packaging plasmid psPAX2 (Addgene; #12260), and 2.5 µg envelope plasmid for the expression of Spike glycoprotein (obtained as a kind gift from Prof. Nevan Krogan, UCSF, USA) of SARS-CoV-2 were mixed in 500 µl OptiMEM media and 20 µl PLUS reagent (Invitrogen; LTX transfection reagent, catalogue no. L15338100) and kept for incubation at room temperature for 5min. In a separate microcentrifuge vial, 500 µl OptiMEM was taken, and 30 µl Lipofectamine-LTX reagent was added to it. This Lipofectamine-containing solution was added to the plasmid and incubated at room temperature for 20 min. We also generated control lentiviral particles by replacing Spike plasmid with VSV-G envelope protein, pmDG2 (Addgene; #12259) plasmid. The above transfection mix was added to the cells and post 16-18hr the media was changed to PenStrep containing complete DMEM media. The viral supernatant was collected at 48hr, 72hr, and 96hr post-transfection by replacing 10ml fresh media every time. All the viral supernatant were pooled together and stored at 4°C till 96hr post-transfection. The supernatant was concentrated up to ∼1–3 ml using a 50-KDa Millipore Amicon filter (Merck; UFC905024) at 1,000*g*, and 4°C. The concentrated viral supernatant was mixed with one-third volume of Lenti-X concentrator (Takara Bio; catalogue no. 631231) overnight at 4°C and pelleted at 1,500*g* for 45 min at 4°C. The off-white pellet of the viruses was resuspended in 1–2 ml of complete DMEM media (10% FBS) and stored at −80°C in 100-200 µl aliquots until further use.

### Omicron pseudotyped virus production

Omicron pseudotyped viruses were produced similarly as described above for spike pseudoviruses but instead used omicron envelope plasmid along with packaging plasmid (psPAX2) and lentiviral plasmid (pHR mCherry) in the following ratio: psPAX2 (1.3pmol), pHR mCherry: 1.64pmol, SARS-CoV-2 Omicron Strain S gene (Genscript, Cat # MC_0101274): 0.72pmol.

### Cloning of ACE2 in lentiviral vector (pTRIP vector)

Human ACE2 was amplified from the mammalian Caco2 cell lines. Caco2 cells were lysed for total RNA purification using the Trizol method. Further, 2 µg of total RNA was set up for cDNA synthesis with Verso cDNA synthesis kit (ThermoFisher Scientific, catalogue no. AB1453A) as per the manufacturer’s protocol. The protein-coding sequence of ACE2 was amplified from the cDNA with forward (5’-ggaggagaaccctggacctggatccatgtcaagctcttcctggctcc-3’) and reverse (5’-ctcctgaccctcctcccccgtaaaaggaggtctgaacatc-3’) primers using Q5 polymerase PCR reaction (NEB, catalogue no. M0492L). This amplified construct of ACE2 was cloned into a pTRIP chicken β-actin (CAG) vector (Gentili et al., 2015). This construct contains amino-terminus EGFP followed by self-cleaving 2A peptide sequence followed by ACE2 and carboxy-termini SNAP-tag and FLAG tags (eGFP-ACE2/HEK293T)

### Generation of stable HEK293T cell line for over-expression of ACE2

The lentiviral pTRIP vector cloned with eGFP-ACE2/HEK293T under CMV enhancer and chicken β-actin promoter (CAG promoter) flanked with 5′and 3′ long terminal repeat (LTR) sequences (39), were used to produce lentiviral particles as per the method described before (35). Briefly, 70-80% confluent HEK293T cells were transfected with 5 µg lentiviral construct of eGFP-ACE2/HEK293Texpression, 3.75 µg psPAX2 (Addgene; #12260), and 2.5 µg pmDG2 (Addgene; #12259) plasmids using Lipofectamine-LTX reagent. The lentiviruses collected at 48hr, 72, and 96hr were concentrated and pelleted using a lenti-X concentrator. The white pellet of lentiviruses was resuspended in 2ml of complete DMEM media and 1ml of this virus was used to transduce HEK293T cells for stable expression of ACE2 in these cells. We could only achieve the transduction efficiency of 60-70% and therefore, we performed fluorescent activated cell sorting (FACS) experiments to enrich the eGFP expressing (a proxy for ACE2 expression) HEK293T cells in the culture. In our sorted culture, >90% of cells were found to be eGFP positive or expressing eGFP-ACE2/HEK293T

### Immunofluorescence assay for ACE2 expression

The stable HEK293T cells were seeded in ibidi glass-bottom dishes (catalogue no. 81218) precoated with poly-D-Lysine and grown to 70% confluency. The cells were washed with 1xBRB80 (80 mM Pipes, 1 mM MgCl_2_, and 1 mM EGTA, pH 6.8) buffer twice and fixed using 100% ice-cold Methanol for 10min at -20°C. Post fixation cells were washed twice with 1xBRB80 buffer and permeabilized in the same buffer having 0.1% Triton-X-100 for 10min at room temperature. Cells were blocked with 5% BSA made in 1xBRB80 + 0.1% Triton-X-100 for 1hr at room temperature. Cells were incubated with 1:500 dilution of mouse anti-FLAG monoclonal antibody (Merck, catalogue no. F3165) overnight at 4°C. Further, cells were washed thrice for 5min using blocking buffer, followed by secondary goat anti-mouse Alexa Fluor-647 antibody (Invitrogen; 1:1000 dilution) incubation in the same blocking buffer at room temperature for 2hr. Cells were stained for nucleus using DAPI (1 μg/ml) for 10 min followed by five washes with 1xBRB80 for 5min. Cells were imaged on an H-TIRF microscope using 405nm, 488nm and 640nm laser with appropriate filter sets.

### Immunoblotting assay for ACE2 expression

The HEK293T cells were seeded in 35mm dish grown to 70% confluency. The cells were transfected with 1µg of eGFP-ACE2/HEK293T with Jet Prime reagent for 24hrs and 48hrs. Cells were lysed with RIPA buffer and subjected to 12% SDS-PAGE. The wet transfer was performed on methanol activated PVDF membrane for 75mins at 100V at 4□C. The blots were probed with 1:5000 anti Flag antibody (Sigma, F3165) overnight at 4□C followed by 1:10000 anti-mouse secondary at room temperature for 1hr. The blots were developed by chemiluminescence on iBright1500 (Invitrogen) by Pierce ECL substrate (Thermo Fisher Scientific; Catalog no. 32106).

### ELISA assay

The surface of the 96 well plates was passivated with BSA-biotin (Thermo Fisher Scientific; catalogue no. 29130, 1 mg/ml, 5 min), followed by a 1×PBS wash. The surface was coated with 0.5 mg/ml Streptavidin (Thermo Fisher Scientific; catalogue no. 43–4302) for 5 min and washed twice with 1xPBS (8 g/liter NaCl, 0.2 g/liter KCl, 1.44 g/liter Na_2_HPO_4_, 0.24 g/liter KH_2_PO_4_, pH 7.4). The surface was immobilised with a 200 μM biotinylated peptide-1/peptide-2 sequence for 30min on ice. The well surface was blocked with 5% BSA containing 1xPBS for 1hr on ice. All the nanobodies (containing carboxy-termini 6x-His-tag) screened for peptide-1 were diluted in blocking buffer at 10 μM concentration and was incubated with the immobilised peptides for 1hr on ice. The surface of the well was washed twice with blocking buffer and incubated with HRP-conjugated rabbit anti-6xHis tag antibody (Abcam; catalogue no. AB1187), 1:10,000 dilution at 4°C overnight. The wells were washed thrice with the blocking buffer and incubated with 100μl of 1:10 diluted (diluted in distilled water) TMB substrate solution (3,3’,5,5’-tetramethylbenzidine; ThermoFisher Scientific, catalogue no. N301). Once the colour starts to develop, stop the reaction using 0.2 mM sulfuric acid. The colour will turn yellow upon sulfuric acid addition, which was measured for absorbance at 405nm using a spectrophotometer keeping appropriate controls (wells not having any immobilised peptides). The intensity of the yellow colour represents the binding of nanobodies with the respective peptide. The relative absorbance (A405) values of individual nanobodies plotted in the graph represent A405 with peptide subtracted from A405 without peptide.

### Pseudoviral transduction assay

The GFP/HEK293T cells or eGFP-ACE2/HEK293T cells were grown up to 60-70% confluency in complete media before viral transduction. In a separate vial mCherry pseudoviruses/ lentiviruses were incubated with the respective purified nanobody at room temperature for 5min. The cells were transduced with the above viral mix along with 2 μg/ml Polybrene (Merck; catalogue no. TR-1003-G) overnight (12-15hr), keeping appropriate positive and negative controls. Post viral incubation, cells were grown in complete media till 48hr and then processed for flow cytometric analysis and confocal imaging.

In this assay, the viral titre was used in a concentration such that to obtain more than 70% transduction efficiency in the eGFP-ACE2/HEK293T or eGFP/HEK293T cell line. We have used 10µl of spike pseudoviruses and 4µl omicron pseudoviruses for flow cytometry and microscopy experiments. For flow cytometric analysis, cells were resuspended in 1xPBS and were subjected to Attune NxT Acoustic Focusing Cytometer to measure red (mCherry) and green (eGFP-ACE2/HEK293T) fluorescence using YL2 (620/15 nm) and BL1 (530/30 nm) filters. Confocal imaging was performed to quantitate the mCherry expression in the stable HEK293T cells after pseudoviral transduction in the presence and absence of the N1.2 and (N1.2)_2_ nanobodies.

Quantification of percentage transduction of cells with pseudovirus (spike and omicron) was done on BD LSRFortessa using 488 and 561 lasers with 530/30 (505LP) and 610/20 (600LP) respectively.

### Imaging and Statistical Analysis

All the images were acquired on an inverted confocal microscope (Olympus FV3000) equipped with six solid state laser lines (405, 445, 488, 514, 561, and 640 nm) and a 20x oil objective. For acquisition and quantification of viral transduction in the pseudoviral assay, high sensitivity spectral detectors were used marking a specific region of interest in the 2048*2048 pixel frame. For all the sets of images acquired for quantification laser power, voltage, and gain settings were kept constant. Images were analysed on Fiji software to calculate mean fluorescence intensity (MFI) for eGFP (ACE2 expression) and mCherry (viral transduction) channel from the z-projected stacks. All the experiments were performed in triplicate sets on at least two different days. Normalised infectivity represents the mean mCherry intensity of a particular z-projected stack over the GFP intensity of the same stacks. Representative images are the z-projected stacks of the respective condition. Respective graphs in Figure-3 and Figure-4 were plotted for the mean intensity and standard error of the mean (s.e.m.)for all the triplicate sets of experiments.

eGFP-ACE2/HEK293T immunostained with FLAG antibody was imaged at 60x oil objective of FV3000 confocal microscope (Figure 2B) and 100x oil objective of H-TIRF microscope (Supplementary Figure 2A) using 405nm, 488nm and 647nm laser lines for DAPI, eGFP, and Alexa fluor-647 fluorophores.

## Acknowledgements

The authors acknowledge the Central Imaging and Flow Facility (CIFF) at the Bangalore Life Science Cluster, India. M.S acknowledges funding support from inStem core grants from the Department of Biotechnology, India, DBT/Wellcome Trust India Alliance Intermediate Fellowship (IA/I/14/2/501533), EMBO Young Investigator Programme award, CEFIPRA (5703-1) from the Department of Science and Technology, SERB-EMR grant (CRG/2019/003246) and DBT-BIRAC (BT/PR40389/COT/142/6/2020) grant. A.C is supported by the DBT/Wellcome Trust India Alliance Early-Career Fellowship (IA/E/15/1/502339). R.G and D.D are supported by a CSIR Fellowship. N.M supported by inStem graduate program.

## Author Contribution

M.S. conceived and supervised the project. N.M, R.G, and S.K screened, validated, and performed the experiments with the nanobody against pseudovirus. D.D, S.S, and P.L. purified the nanobodies used in the ELISA assay. A.C provided inputs in omicron pseudotyped virus production and assays. N.M, R.G, S.K, and M.S interpreted the results and wrote the manuscript and all authors commented on the manuscript content.

---

**Supplementary Figure 1:**
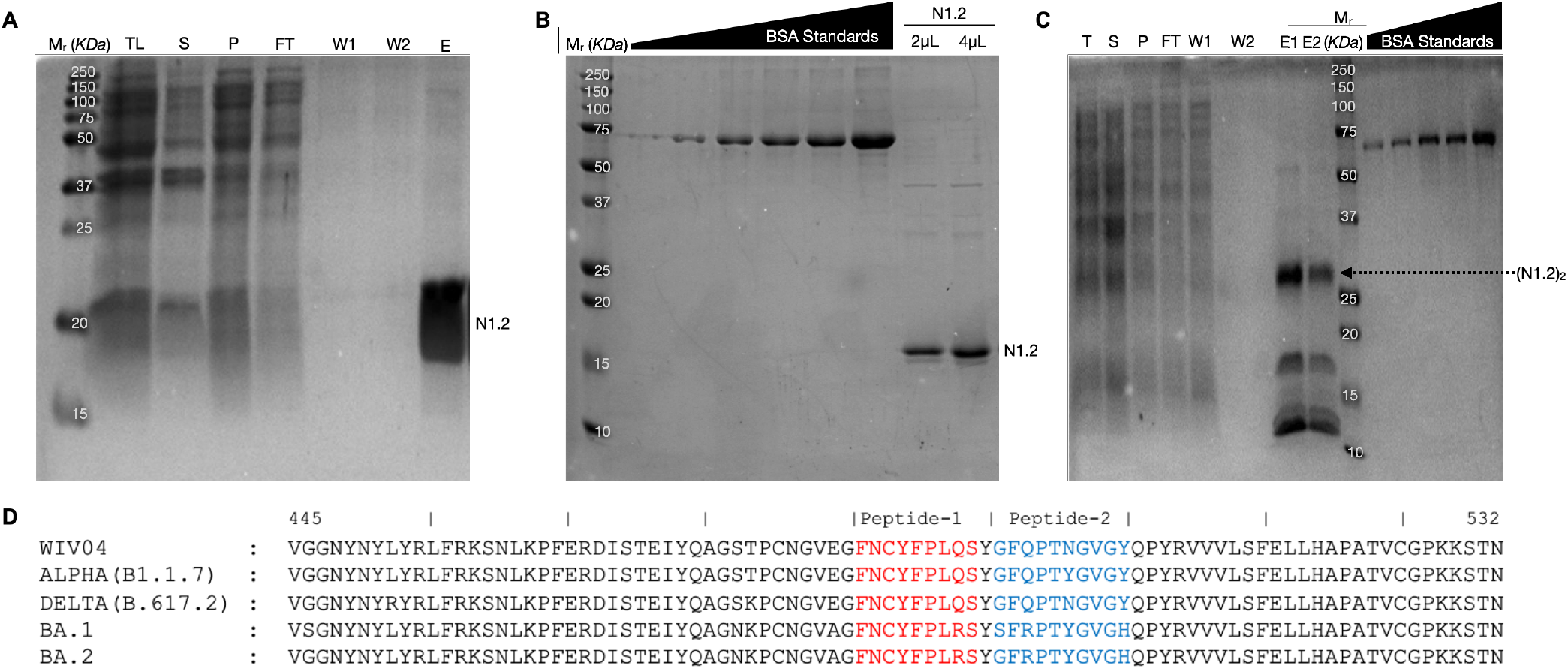
Establishment of virus titer for cell assay. **A, B, C** SDS-PAGE gel of nanobody N1.2 and (N1.2)_2_ was used in this study. **D**. Sequence alignment of RBM (445-532) region of WIV04, Alpha, Delta, and Omicron variants.

**Supplementary Figure 2:**
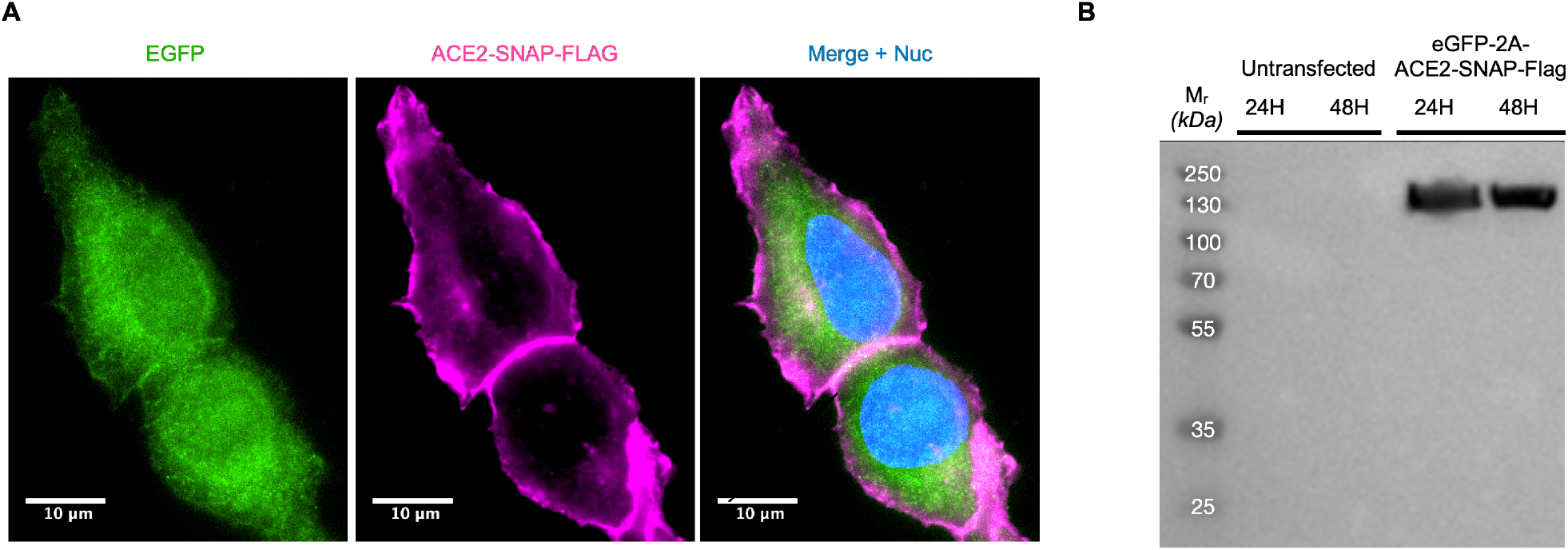
Probing the eGFP-ACE2/HEK293T stable cell line. **A**. TIRF images of eGFP-ACE2/HEK293T cell line developed in this study (Methods). In the DNA sequence, the eGFP and ACE2-SNAP-FLAG genes are separated by the P2A site resulting in two separate protein entities. The eGFP fluorescence (in green) indicates the expression of both eGFP and ACE2-SNAP-FLAG fusion (in magenta) and the nucleus is shown in blue. The ACE2-SNAP-FLAG fusion protein was probed using an anti-FLAG antibody (Methods). **B**. Western blot analysis of eGFP-ACE2/HEK293T stable cell lysate probes using anti-FLAG antibody, the lanes and molecular weight marker are indicated.

**Supplementary Figure 3:**
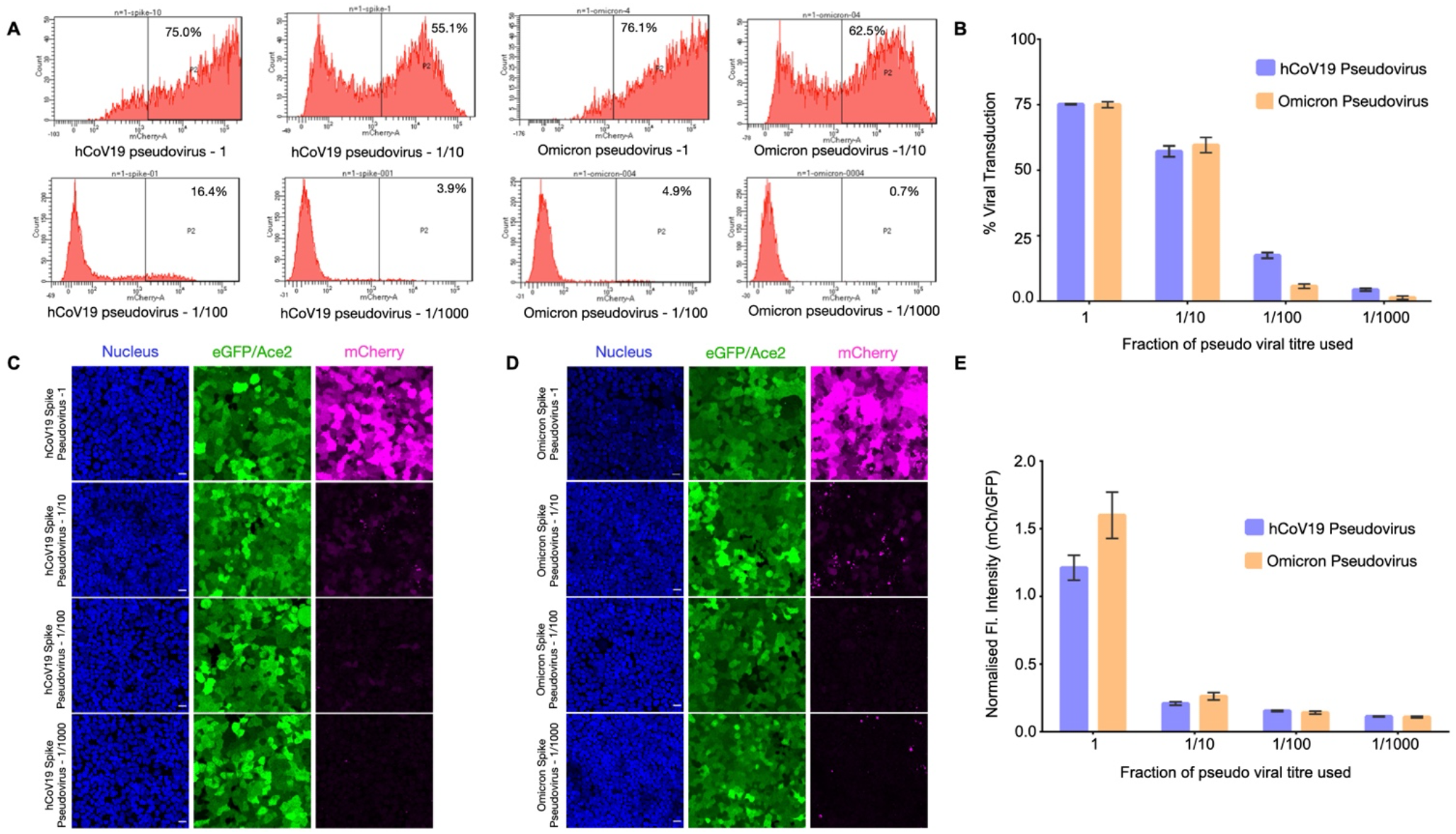
Establishment of virus titer for cell assay. **A**. Raw FACS plots of mCherry positive cells for various dilution of virus used in infecting the eGFP-ACE2/HEK293T cells as indicated. **B**. Percentage of viral transduction for WIV04 (in blue) and omicron (in yellow) pseudovirus quantified from the percentage of mCherry positive cells using FACS analysis. The dilutions for hCoV19 pseudovirus are 10ul, 1ul, 0.1ul, and 0.01ul and omicron pseudovirus are 4ul, 0.4ul, 0.04ul, and 0.004ul. **C and D**. Maximum intensity Z-projection of confocal images of hCoV19and omicron spike pseudovirus for various virus dilutions as indicated. The nucleus is indicated in blue, eGFP in green, and mCherry (reporter) in magenta. Scale bar = 20 microns. **E**. Normalized mCherry/eGFP fluorescence intensity for WIV04 (in blue) and omicron (in yellow) pseudovirus for various virus dilutions as indicated.

**Supplementary Figure 4:**
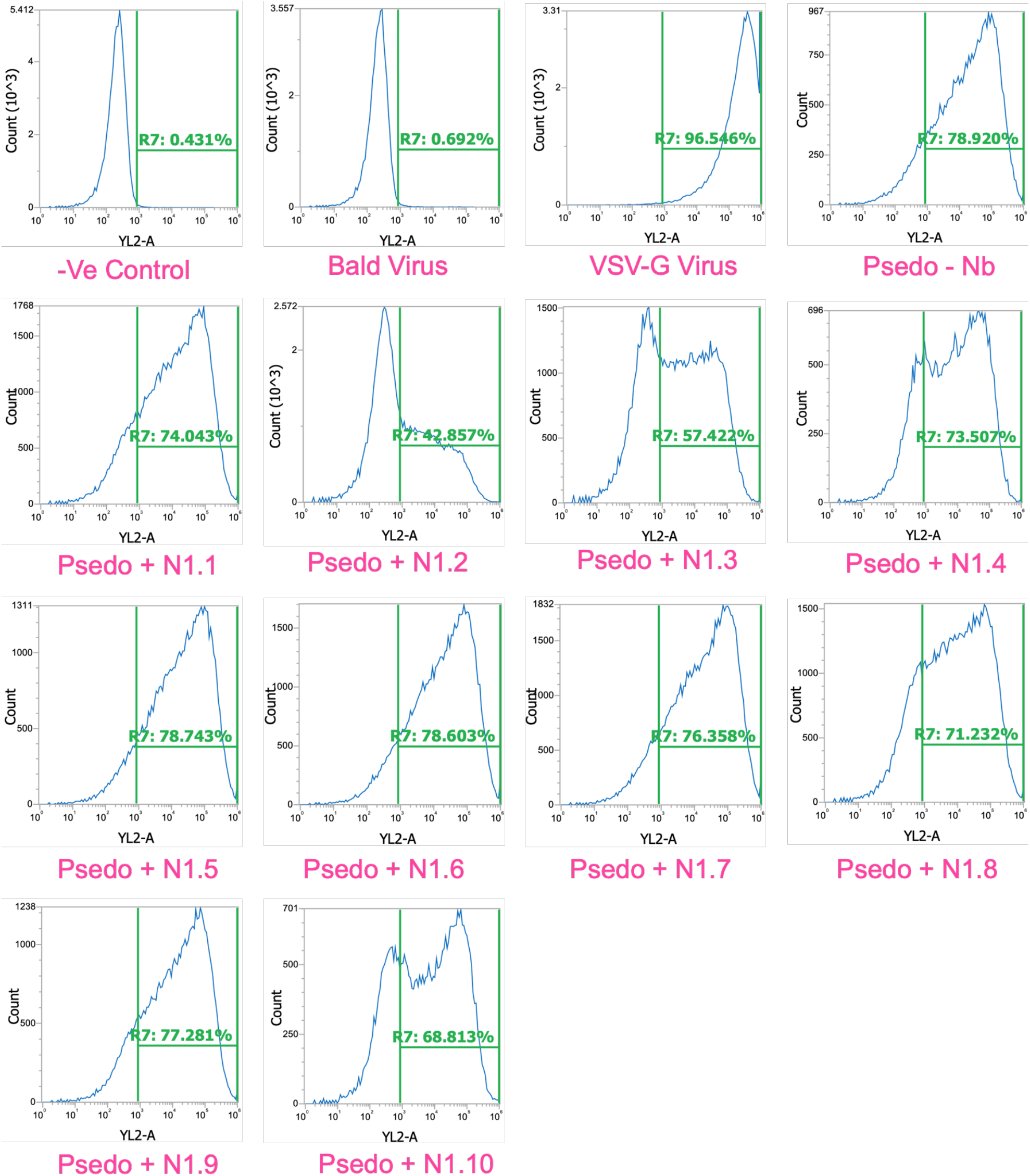
FACS plots of pseudovirus assay. **A and B**. Raw FACS plots used in Figure 2C quantification of the percentage of cells that are positive with mCherry fluorescence after infection with pseudovirus in the presence and absence of nanobodies as indicated. The green vertical line indicates the detection cutoff used for mCherry fluorescence.

**Supplementary Figure 5:**
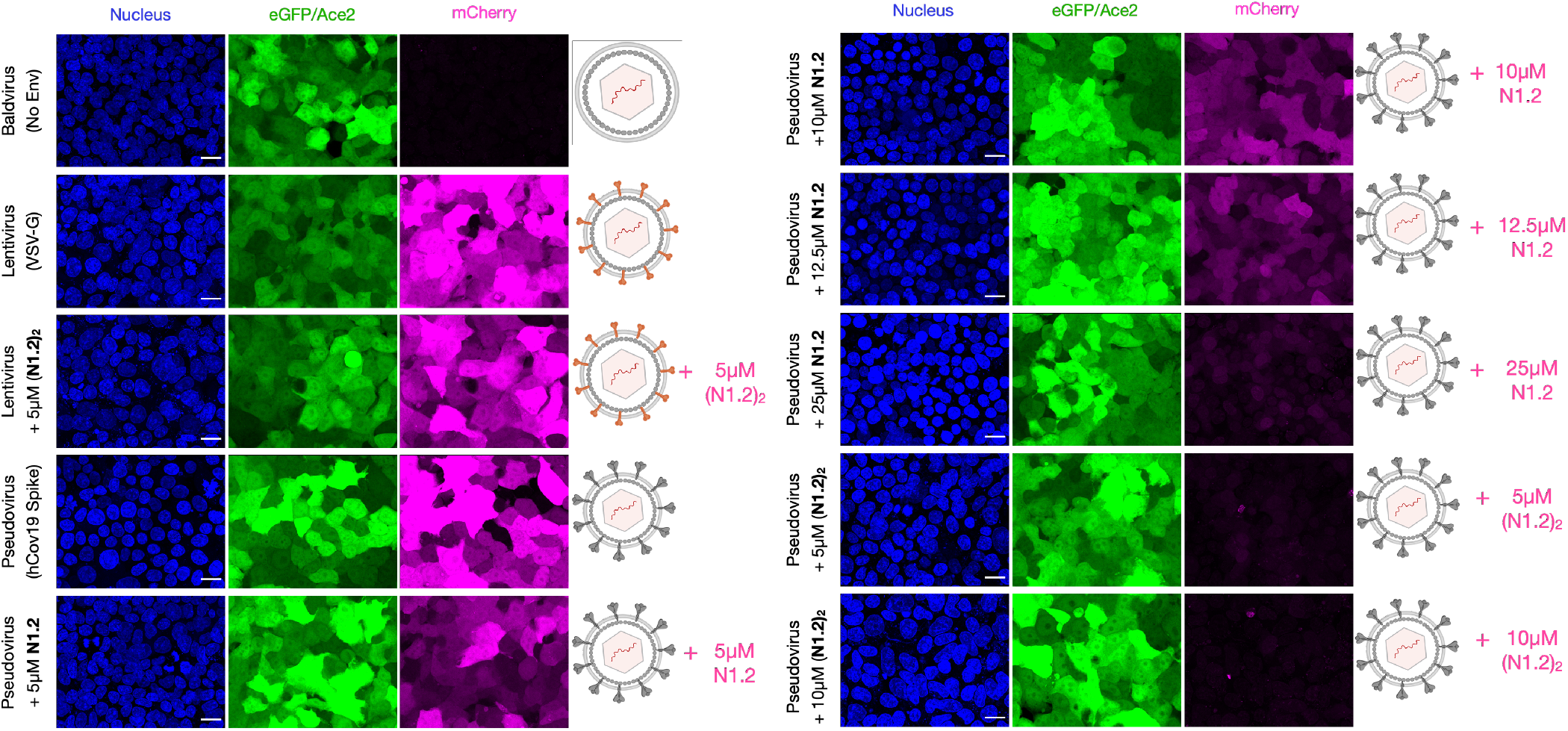
Confocal images of pseudovirus assay against N1.2. Maximum intensity Z-projection of confocal images of eGFP-ACE2/HEK293T cells with bald, VSV-G, and spike pseudovirus in the absence and presence of various concentrations of monovalent and bivalent nanobody N1.2 as indicated. The nucleus is shown in blue, the eGFP fluorescence (in green), and mCherry (in magenta) indicates the extent of virus entry into the ACE2-HEK293T cells. Scale bar = 20 microns.

**Supplementary Figure 6:**
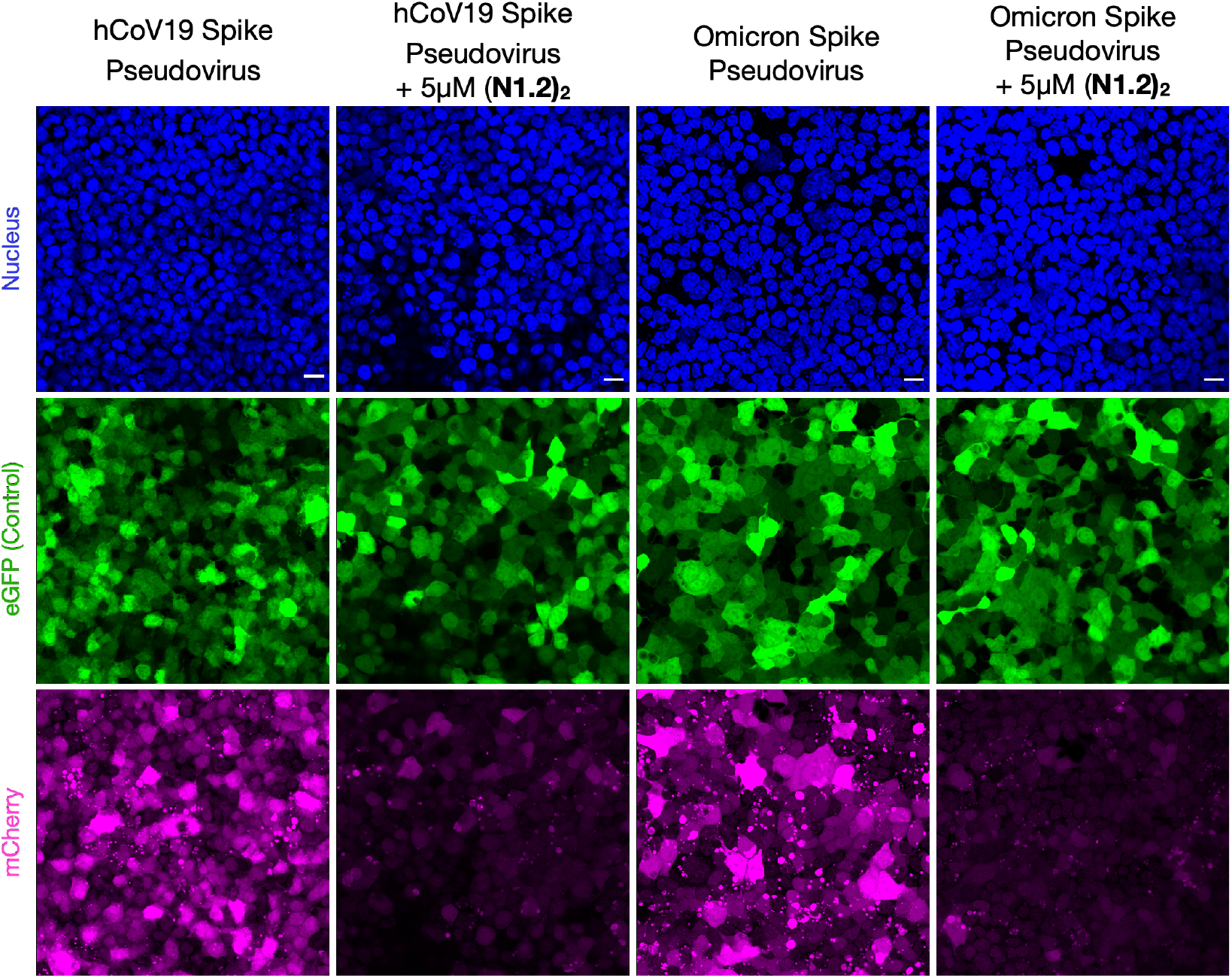
Confocal images of pseudovirus assay with HEK293T cells. Maximum intensity Z-projection of confocal images of GFP/HEK293T cells with WIV04 and Omicron spike pseudovirus against 5uM bivalent nanobody N1.2 as indicated. The nucleus is shown in blue, the eGFP fluorescence (in green) and mCherry (in magenta) indicates the extent of virus entry into the HEK293T cells. Scale bar = 20 microns.

**Supplementary Table 1:**
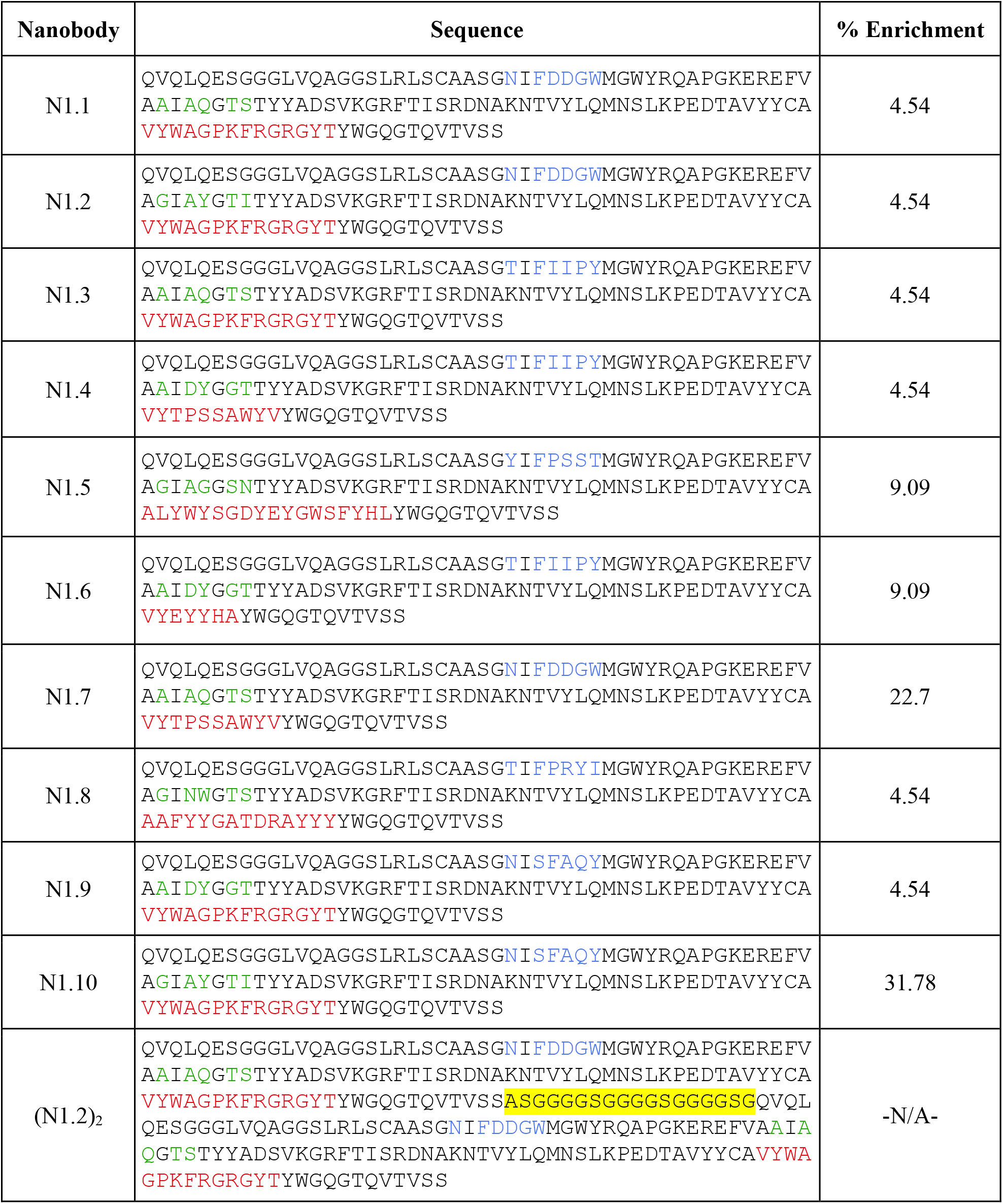
Nanobody clone analysis. Sequences of nanobodies analyzed after enrichment, the three variable regions of the VHH sequence are highlighted in blue, green, and red colors. The percentage of enrichment indicates the reoccurrence of a particular nanobody sequence from 22 yeast colonies isolated.

## References

1. Tang, D., Comish, P., and Kang, R. (2020) The hallmarks of COVID-19 disease. PLOS Pathog. 16, e1008536

2. Xu, Z., Liu, K., and Gao, G. F. (2022) Omicron variant of SARS-CoV-2 imposes a new challenge for the global public health. Biosaf. Health. 10.1016/j.bsheal.2022.01.002

3. Cobey, S., Larremore, D. B., Grad, Y. H., and Lipsitch, M. (2021) Concerns about SARS-CoV-2 evolution should not hold back efforts to expand vaccination. Nat. Rev. Immunol. 21, 330–335

4. He, Y., Lu, H., Siddiqui, P., Zhou, Y., and Jiang, S. (2005) Receptor-binding domain of severe acute respiratory syndrome coronavirus spike protein contains multiple conformation-dependent epitopes that induce highly potent neutralizing antibodies. J. Immunol. Baltim. Md 1950. 174, 4908–4915

5. Lan, J., Ge, J., Yu, J., Shan, S., Zhou, H., Fan, S., Zhang, Q., Shi, X., Wang, Q., Zhang, L., and Wang, X. (2020) Structure of the SARS-CoV-2 spike receptor-binding domain bound to the ACE2 receptor. Nature. 10.1038/s41586-020-2180-5

6. Walls, A. C., Park, Y.-J., Tortorici, M. A., Wall, A., McGuire, A. T., and Veesler, D. (2020) Structure, Function, and Antigenicity of the SARS-CoV-2 Spike Glycoprotein. Cell. 181, 281-292.e6

7. Shang, J., Wan, Y., Luo, C., Ye, G., Geng, Q., Auerbach, A., and Li, F. (2020) Cell entry mechanisms of SARS-CoV-2. Proc. Natl. Acad. Sci. U. S. A. 117, 11727–11734

8. Custódio, T. F., Das, H., Sheward, D. J., Hanke, L., Pazicky, S., Pieprzyk, J., Sorgenfrei, M., Schroer, M. A., Gruzinov, A. Y., Jeffries, C. M., Graewert, M. A., Svergun, D. I., Dobrev, N., Remans, K., Seeger, M. A., McInerney, G. M., Murrell, B., Hällberg, B. M., and Löw, C. (2020) Selection, biophysical and structural analysis of synthetic nanobodies that effectively neutralize SARS-CoV-2. Nat. Commun. 11, 5588

9. Hanke, L., Vidakovics Perez, L., Sheward, D. J., Das, H., Schulte, T., Moliner-Morro, A., Corcoran, M., Achour, A., Karlsson Hedestam, G. B., Hällberg, B. M., Murrell, B., and McInerney, G. M. (2020) An alpaca nanobody neutralizes SARS-CoV-2 by blocking receptor interaction. Nat. Commun. 11, 4420

10. Huo, J., Bas, A. L., Ruza, R., Duyvesteyn, H. M. E., Mikolajek, H., Malinauskas, T., Tan, T. K., Rijal, P., Dumoux, M., Ward, P., Ren, J., Zhou, D., Harrison, P., Radeke, J., Zhao, Y., Gilbert-Jaramillo, J., Knight, M., Carrique, L., Shah, P. N. M., James, W., Townsend, A., Stuart, D., Owens, R., and Naismith, J. (2020) Structural characterisation of a nanobody derived from a naïve library that neutralises SARS-CoV-2. 10.21203/rs.3.rs-32948/v1

11. Mast, F. D., Fridy, P. C., Ketaren, N. E., Wang, J., Jacobs, E. Y., Olivier, J. P., Sanyal, T., Molloy, K. R., Schmidt, F., Rutkowska, M., Weisblum, Y., Rich, L. M., Vanderwall, E. R., Dambrauskas, N., Vigdorovich, V., Keegan, S., Jiler, J. B., Stein, M. E., Olinares, P. D. B., Herlands, L., Hatziioannou, T., Sather, D. N., Debley, J. S., Fenyö, D., Sali, A., Bieniasz, P. D., Aitchison, J. D., Chait, B. T., and Rout, M. P. (2021) Highly synergistic combinations of nanobodies that target SARS-CoV-2 and are resistant to escape. eLife. 10, e73027

12. Shi, R., Shan, C., Duan, X., Chen, Z., Liu, P., Song, J., Song, T., Bi, X., Han, C., Wu, L., Gao, G., Hu, X., Zhang, Y., Tong, Z., Huang, W., Liu, W. J., Wu, G., Zhang, B., Wang, L., Qi, J., Feng, H., Wang, F.-S., Wang, Q., Gao, G. F., Yuan, Z., and Yan, J. (2020) A human neutralizing antibody targets the receptor-binding site of SARS-CoV-2. Nature. 584, 120–124

13. Wang, C., Li, W., Drabek, D., Okba, N. M. A., van Haperen, R., Osterhaus, A. D. M. E., van Kuppeveld, F. J. M., Haagmans, B. L., Grosveld, F., and Bosch, B.-J. (2020) A human monoclonal antibody blocking SARS-CoV-2 infection. Nat. Commun. 11, 2251

14. Wrapp, D., De Vlieger, D., Corbett, K. S., Torres, G. M., Wang, N., Van Breedam, W., Roose, K., van Schie, L., VIB-CMB COVID-19 Response Team, Hoffmann, M., Pöhlmann, S., Graham, B. S., Callewaert, N., Schepens, B., Saelens, X., and McLellan, J. S. (2020) Structural Basis for Potent Neutralization of Betacoronaviruses by Single-Domain Camelid Antibodies. Cell. 181, 1436–1441

15. Xiang, Y., Nambulli, S., Xiao, Z., Liu, H., Sang, Z., Duprex, W. P., Schneidman-Duhovny, D., Zhang, C., and Shi, Y. (2020) Versatile and multivalent nanobodies efficiently neutralize SARS-CoV-2. Science. 370, 1479–1484

16. Ye, M., Fu, D., Ren, Y., Wang, F., Wang, D., Zhang, F., Xia, X., and Lv, T. (2020) Treatment with convalescent plasma for COVID-19 patients in Wuhan, China. J. Med. Virol. 92, 1890–1901

17. B, Z., S, L., T, T., W, H., Y, D., L, C., Q, C., L, Z., Q, Z., X, Z., Y, Z., and S, Z. (2020) Treatment With Convalescent Plasma for Critically Ill Patients With Severe Acute Respiratory Syndrome Coronavirus 2 Infection. Chest. 10.1016/j.chest.2020.03.039

18. Cao, Y., Su, B., Guo, X., Sun, W., Deng, Y., Bao, L., Zhu, Q., Zhang, X., Zheng, Y., Geng, C., Chai, X., He, R., Li, X., Lv, Q., Zhu, H., Deng, W., Xu, Y., Wang, Y., Qiao, L., Tan, Y., Song, L., Wang, G., Du, X., Gao, N., Liu, J., Xiao, J., Su, X.-D., Du, Z., Feng, Y., Qin, C., Qin, C., Jin, R., and Xie, X. S. (2020) Potent Neutralizing Antibodies against SARS-CoV-2 Identified by High-Throughput Single-Cell Sequencing of Convalescent Patients’ B Cells. Cell. 182, 73-84.e16

19. Minenkova, O., Santapaola, D., Milazzo, F. M., Anastasi, A. M., Battistuzzi, G., Chiapparino, C., Rosi, A., Gritti, G., Borleri, G., Rambaldi, A., Dental, C., Viollet, C., Pagano, B., Salvini, L., Marra, E., Luberto, L., Rossi, A., Riccio, A., Pich, E. M., Santoro, M. G., and Santis, R. D. (2022) Human inhalable antibody fragments neutralizing SARS-CoV-2 variants for COVID-19 therapy. Mol. Ther. 10.1016/j.ymthe.2022.02.013

20. Shen, C., Wang, Z., Zhao, F., Yang, Y., Li, J., Yuan, J., Wang, F., Li, D., Yang, M., Xing, L., Wei, J., Xiao, H., Yang, Y., Qu, J., Qing, L., Chen, L., Xu, Z., Peng, L., Li, Y., Zheng, H., Chen, F., Huang, K., Jiang, Y., Liu, D., Zhang, Z., Liu, Y., and Liu, L. (2020) Treatment of 5 Critically Ill Patients With COVID-19 With Convalescent Plasma. JAMA. 323, 1582–1589

21. Wu, X., Cheng, L., Fu, M., Huang, B., Zhu, L., Xu, S., Shi, H., Zhang, D., Yuan, H., Nawaz, W., Yang, P., Hu, Q., Liu, Y., and Wu, Z. (2021) A potent bispecific nanobody protects hACE2 mice against SARS-CoV-2 infection via intranasal administration. Cell Rep. 37, 109869

22. Ye, G., Gallant, J., Zheng, J., Massey, C., Shi, K., Tai, W., Odle, A., Vickers, M., Shang, J., Wan, Y., Du, L., Aihara, H., Perlman, S., LeBeau, A., and Li, F. (2021) The development of Nanosota-1 as anti-SARS-CoV-2 nanobody drug candidates. eLife. 10, e64815

23. Muyldermans, S. (2013) Nanobodies: natural single-domain antibodies. Annu. Rev. Biochem. 82, 775–797

24. Bannas, P., Hambach, J., and Koch-Nolte, F. (2017) Nanobodies and Nanobody-Based Human Heavy Chain Antibodies As Antitumor Therapeutics. Front. Immunol. 8, 1603

25. Vincke, C., Loris, R., Saerens, D., Martinez-Rodriguez, S., Muyldermans, S., and Conrath, K. (2009) General strategy to humanize a camelid single-domain antibody and identification of a universal humanized nanobody scaffold. J. Biol. Chem. 284, 3273– 3284

26. McMahon, C., Baier, A. S., Pascolutti, R., Wegrecki, M., Zheng, S., Ong, J. X., Erlandson, S. C., Hilger, D., Rasmussen, S. G. F., Ring, A. M., Manglik, A., and Kruse, A. C. (2018) Yeast surface display platform for rapid discovery of conformationally selective nanobodies. Nat. Struct. Mol. Biol. 25, 289–296

27. Chen, M., and Zhang, X.-E. (2021) Construction and applications of SARS-CoV-2 pseudoviruses: a mini review. Int. J. Biol. Sci. 17, 1574–1580

28. Hoffmann, M., Kleine-Weber, H., Schroeder, S., Krüger, N., Herrler, T., Erichsen, S., Schiergens, T. S., Herrler, G., Wu, N.-H., Nitsche, A., Müller, M. A., Drosten, C., and Pöhlmann, S. (2020) SARS-CoV-2 Cell Entry Depends on ACE2 and TMPRSS2 and Is Blocked by a Clinically Proven Protease Inhibitor. Cell. 181, 271-280.e8

29. Sherman, E. J., and Emmer, B. T. (2021) ACE2 protein expression within isogenic cell lines is heterogeneous and associated with distinct transcriptomes. bioRxiv. 10.1101/2021.03.26.437218

30. Harvey, W. T., Carabelli, A. M., Jackson, B., Gupta, R. K., Thomson, E. C., Harrison, E. M., Ludden, C., Reeve, R., Rambaut, A., COVID-19 Genomics UK (COG-UK) Consortium, Peacock, S. J., and Robertson, D. L. (2021) SARS-CoV-2 variants, spike mutations and immune escape. Nat. Rev. Microbiol. 19, 409–424

31. Du, L., Yang, Y., and Zhang, X. (2021) Neutralizing antibodies for the prevention and treatment of COVID-19. Cell. Mol. Immunol. 18, 2293–2306

32. Papageorgiou, A. C., and Mohsin, I. (2020) The SARS-CoV-2 Spike Glycoprotein as a Drug and Vaccine Target: Structural Insights into Its Complexes with ACE2 and Antibodies. Cells. 9, E2343

33. Yu, F., Xiang, R., Deng, X., Wang, L., Yu, Z., Tian, S., Liang, R., Li, Y., Ying, T., and Jiang, S. (2020) Receptor-binding domain-specific human neutralizing monoclonal antibodies against SARS-CoV and SARS-CoV-2. Signal Transduct. Target. Ther. 5, 212

34. Traboulsi, H., Khedr, M. A., Al-Faiyz, Y. S. S., Elgorashe, R., and Negm, A. (2021) Structure-Based Epitope Design: Toward a Greater Antibody–SARS-CoV-2 RBD Affinity. ACS Omega. 6, 31469–31476

35. Kesarwani, S., Lama, P., Chandra, A., Reddy, P. P., Jijumon, A. S., Bodakuntla, S., Rao, B. M., Janke, C., Das, R., and Sirajuddin, M. (2020) Genetically encoded live-cell sensor for tyrosinated microtubules. J. Cell Biol. 219, e201912107

36. Sanches, P. R. S., Charlie-Silva, I., Braz, H. L. B., Bittar, C., Freitas Calmon, M., Rahal, P., and Cilli, E. M. (2021) Recent advances in SARS-CoV-2 Spike protein and RBD mutations comparison between new variants Alpha (B.1.1.7, United Kingdom), Beta (B.1.351, South Africa), Gamma (P.1, Brazil) and Delta (B.1.617.2, India). J. Virus Erad. 7, 100054

37. Wu, L., Zhou, L., Mo, M., Liu, T., Wu, C., Gong, C., Lu, K., Gong, L., Zhu, W., and Xu, Z. (2022) SARS-CoV-2 Omicron RBD shows weaker binding affinity than the currently dominant Delta variant to human ACE2. Signal Transduct. Target. Ther. 7, 8

38. Gordon, D. E., Jang, G. M., Bouhaddou, M., Xu, J., Obernier, K., White, K. M., O’Meara, M. J., Rezelj, V. V., Guo, J. Z., Swaney, D. L., Tummino, T. A., Hüttenhain, R., Kaake, R. M., Richards, A. L., Tutuncuoglu, B., Foussard, H., Batra, J., Haas, K., Modak, M., Kim, M., Haas, P., Polacco, B. J., Braberg, H., Fabius, J. M., Eckhardt, M., Soucheray, M., Bennett, M. J., Cakir, M., McGregor, M. J., Li, Q., Meyer, B., Roesch, F., Vallet, T., Mac Kain, A., Miorin, L., Moreno, E., Naing, Z. Z. C., Zhou, Y., Peng, S., Shi, Y., Zhang, Z., Shen, W., Kirby, I. T., Melnyk, J. E., Chorba, J. S., Lou, K., Dai, S. A., Barrio-Hernandez, I., Memon, D., Hernandez-Armenta, C., Lyu, J., Mathy, C. J. P., Perica, T., Pilla, K. B., Ganesan, S. J., Saltzberg, D. J., Rakesh, R., Liu, X., Rosenthal, S. B., Calviello, L., Venkataramanan, S., Liboy-Lugo, J., Lin, Y., Huang, X.-P., Liu, Y., Wankowicz, S. A., Bohn, M., Safari, M., Ugur, F. S., Koh, C., Savar, N. S., Tran, Q. D., Shengjuler, D., Fletcher, S. J., O’Neal, M. C., Cai, Y., Chang, J. C. J., Broadhurst, D. J., Klippsten, S., Sharp, P. P., Wenzell, N. A., Kuzuoglu-Ozturk, D., Wang, H.-Y., Trenker, R., Young, J. M., Cavero, D. A., Hiatt, J., Roth, T. L., Rathore, U., Subramanian, A., Noack, J., Hubert, M., Stroud, R. M., Frankel, A. D., Rosenberg, O. S., Verba, K. A., Agard, D. A., Ott, M., Emerman, M., Jura, N., von Zastrow, M., Verdin, E., Ashworth, A., Schwartz, O., d’Enfert, C., Mukherjee, S., Jacobson, M., Malik, H. S., Fujimori, D. G., Ideker, T., Craik, C. S., Floor, S. N., Fraser, J. S., Gross, J. D., Sali, A., Roth, B. L., Ruggero, D., Taunton, J., Kortemme, T., Beltrao, P., Vignuzzi, M., García-Sastre, A., Shokat, K. M., Shoichet, B. K., and Krogan, N. J. (2020) A SARS-CoV-2 protein interaction map reveals targets for drug repurposing. Nature. 583, 459–468

39. Gentili, M., Kowal, J., Tkach, M., Satoh, T., Lahaye, X., Conrad, C., Boyron, M., Lombard, B., Durand, S., Kroemer, G., Loew, D., Dalod, M., Théry, C., and Manel, N. (2015) Transmission of innate immune signaling by packaging of cGAMP in viral particles. Science. 349, 1232–1236

